# Quantitative methods to investigate the 4D dynamics of heterochromatic repair sites in Drosophila cells

**DOI:** 10.1101/214601

**Authors:** Christopher P. Caridi, Laetitia Delabaere, Harianto Tjong, Hannah Hopp, Devika Das, Frank Alber, Irene Chiolo

## Abstract

Heterochromatin is mostly composed of long stretches of repeated DNA sequences prone to ectopic recombination during double-strand break (DSB) repair. In *Drosophila*, ‘safe’ homologous recombination (HR) repair of heterochromatic DSBs relies on a striking relocalization of repair sites to the nuclear periphery. Central to understanding heterochromatin repair is the ability to investigate the 4D dynamics (movement in space and time) of repair sites. A specific challenge of these studies is preventing phototoxicity and photobleaching effects while imaging the sample over long periods of time, and with sufficient time points and Z-stacks to track repair foci over time. Here we describe an optimized approach for high-resolution live imaging of heterochromatic DSBs in *Drosophila* cells, with a specific emphasis on the fluorescent markers and imaging setup used to capture the motion of repair foci over long time periods. We detail approaches that minimize photobleaching and phototoxicity with a DeltaVision widefield deconvolution microscope, and image-processing techniques for signal recovery post-imaging using SoftWorX and Imaris software. We present a method to derive mean square displacement (MSD) curves revealing some of the biophysical properties of the motion. Finally, describe a method in R to identify tracts of directed motions in mixed trajectories. These approaches enable a deeper understanding of the mechanisms of heterochromatin dynamics and genome stability in the three-dimensional context of the nucleus, and have broad applicability in the field of nuclear dynamics.

## Nuclear dynamics play critical roles in heterochromatin repair and genome stability

Double strand breaks (DSBs) in pericentromeric heterochromatin (hereafter, ‘heterochromatin’) are a major threat to genome stability (recently reviewed in (Amaral, Ryu, Li, & Chiolo, 2017; Caridi, Delabaere, Zapotoczny, & Chiolo, 2017; Chiolo, Tang, Georgescu, & Costes, 2013)). Heterochromatin comprises ~30% of fly and human genomes (Ho et al., 2014; Roger A. Hoskins et al., 2007; R. A. Hoskins et al., 2015) (Figure 1A) and is mostly composed of repeated sequences prone to ectopic recombination during DNA repair (Amaral et al., 2017; P. C. Caridi et al., 2017; Chiolo et al., 2013; Peng & Karpen, 2008). In *Drosophila* for example, about half of these sequences consist of simple ‘satellite’ repeats (mostly tandem 5-base pair sequences) repeated for hundreds of kilobases to megabases, while the rest are mostly composed of scrambled clusters of transposable elements and about 250 isolated genes (Ho et al., 2014; Roger A. Hoskins et al., 2007; R. A. Hoskins et al., 2015). While heterochromatin is a major component of the genome in multi-cellular eukaryotes, it is absent in budding yeast.

**Figure 1:**
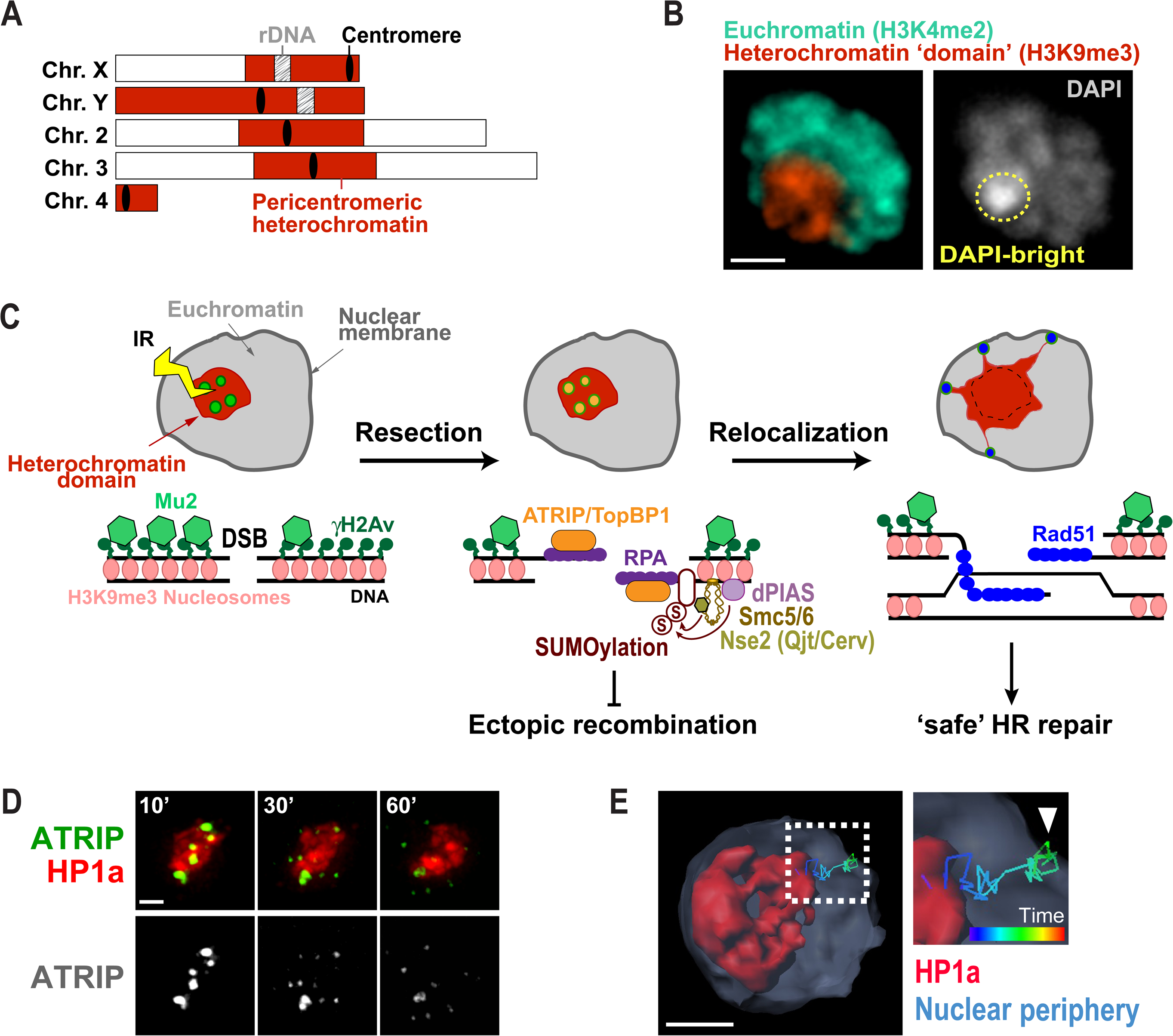
Nuclear dynamics of heterochromatin repair mechanisms in *Drosophila* cells. A) Schematic view of *Drosophila* chromosomes showing the position and extent of pericentromeric heterochromatin. B) Immunofluorescence of a Kc cell showing heterochromatin organized as one distinct domain (red, H3K9me3) comprising the DAPI bright region (yellow circle). Euchromatin surrounds the heterochromatin domain (green, H3K4me2). Scale bar = 1µm. C) HR repair of heterochromatic DSBs starts inside the domain with H2Av phosphorylation and Mu2/Mdc1 recruitment. Resection also occur inside the domain resulting in ATRIP and TopBP1 foci. Next, the heterochromatin domain expands and repair foci relocalize to the nuclear periphery to form Rad51 foci and continue HR. dPIAS, Nse2/Qjt and Nse2/Cerv SUMO-ligases are required to block HR progression in the domain and prevent ectopic recombination. D) Live imaging of one Kc cell expressing mCh-HP1a and EGFP-ATRIP shows ATRIP foci inside the heterochromatin domain at 10 min after IR, which leave the domain by 60 min after IR (Chiolo et al., 2011). E) 4D reconstruction of one nucleus of a Kc cell expressing mCh-HP1a, mCh-LaminC and GFP-ATRIP, and exposed to IR, shows an example of a heterochromatic repair focus that leaves the heterochromatin domain and reaches the nuclear periphery (Ryu et al., 2015).

Despite the risk of aberrant recombination, homologous recombination (HR) is largely utilized for repairing heterochromatic DSBs in both flies (Chiolo et al., 2011; Janssen et al., 2016; Ryu, Bonner, & Chiolo, 2016; Ryu et al., 2015) and mammalian cells (Beucher et al., 2009; Tsouroula et al., 2016), and studies from our lab and others identified specialized pathways that promote heterochromatin repair while preventing aberrant recombination (reviewed in (Amaral et al., 2017; P. C. Caridi et al., 2017; Chiolo et al., 2013)). Working with *Drosophila* cells, we discovered that ‘safe’ HR repair relies on a striking relocalization of heterochromatic repair sites to the nuclear periphery (C. Caridi et al., 2017; Chiolo et al., 2011; Ryu et al., 2016; Ryu et al., 2015). Repair starts inside the heterochromatin domain – a distinct nuclear structure in flies (Chiolo et al., 2011; Riddle et al., 2011; Ryu et al., 2016; Ryu et al., 2015) (Figure 1B) – and is temporarily halted after resection and ATRIP/TopBP1 focus formation (Chiolo et al., 2011; Ryu et al., 2016; Ryu et al., 2015) (Figure 1C). This block to HR progression is mediated by SUMOylation *via* the SUMO E3 ligases dPIAS and Smc5/6 subunits Nse2/Qjt and Nse2/Cerv (Chiolo et al., 2011; Ryu et al., 2016; Ryu et al., 2015) (Figure 1C). Next, the heterochromatin domain expands during relocalization (Chiolo et al., 2011). This likely reflects a global relaxation of the chromatin and facilitates damage signaling and/or nuclear dynamics. Repair continues with Rad51 recruitment at the nuclear periphery (Chiolo et al., 2011; Ryu et al., 2015) (Figure 1D), revealing a tight regulation of repair progression in space and time. Inactivating the relocalization pathway leads to aberrant recombination between heterochromatic sequences, chromosomal aberrations and heterochromatin instability (C. Caridi et al., 2017; Chiolo et al., 2011; Ryu et al., 2016; Ryu et al., 2015), revealing the importance of these dynamics for genome integrity. In mouse cells, where heterochromatin is organized in several ‘chromocenters’, these domains expand during repair (Ayoub, Jeyasekharan, Bernal, & Venkitaraman, 2008; Burgess, Burman, Kruhlak, & Misteli, 2014; Tsouroula et al., 2016) and repair sites leave the domains before Rad51 recruitment and HR progression (Chiolo et al., 2013; Jakob et al., 2011; Tsouroula et al., 2016). These observations suggest highly conserved strategies for ‘safe’ heterochromatin repair. Relocalization likely prevents ectopic recombination by isolating the damaged site and its homologous sequences (on the homologous chromosome or the sister chromatid) away from similar sequences on ectopic chromosomes, before strand invasion (Amaral et al., 2017; P. C. Caridi et al., 2017; Chiolo et al., 2011; Chiolo et al., 2013; Ryu et al., 2016; Ryu et al., 2015).

In addition, a major body of work in yeast and mammalian cells revealed the dynamic nature of chromatin in both damaged and undamaged regions (reviewed in (Amaral et al., 2017; P. C. Caridi et al., 2017)). Repair sites are highly dynamic during homology search for HR repair (Cho, Dilley, Lampson, & Greenberg, 2014; Vincent Dion, Kalck, Horigome, Towbin, & Gasser, 2012; Mine-Hattab & Rothstein, 2012; Neumann et al., 2012; Saad et al., 2014) or during relocalization of relatively rare classes of DSBs to specific subnuclear compartments. For example, DSBs induced in rDNA of yeast and human cells leave the nucleolus during repair (Torres-Rosell et al., 2007; van Sluis & McStay, 2015). Further, persistent DSBs, eroded telomeres, subtelomeric breaks, and collapsed forks relocalize to the nuclear periphery for repair (Chung et al., 2015; Churikov et al., 2016; Horigome et al., 2016; Horigome et al., 2014; Kalocsay, Hiller, & Jentsch, 2009; Khadaroo et al., 2009; Nagai et al., 2008; Oza, Jaspersen, Miele, Dekker, & Peterson, 2009; Su, Dion, Gasser, & Freudenreich, 2015; Swartz, Rodriguez, & King, 2014; Therizols et al., 2006) (recently reviewed in (Amaral et al., 2017; P. C. Caridi et al., 2017)). Interestingly, SUMOylation appears to drive the relocalization of repair sites in different contexts, revealing conserved pathways for the regulation of nuclear dynamics during DSB repair (Churikov et al., 2016; Horigome et al., 2016; Kalocsay et al., 2009; Khadaroo et al., 2009; Nagai et al., 2008; Oza et al., 2009; Su et al., 2015; Torres-Rosell et al., 2007). Structural components in the cytoplasm appear to influence nuclear dynamics via SUN-KASH protein complexes traversing the nuclear membrane (Lottersberger, Karssemeijer, Dimitrova, & de Lange, 2015; Spichal et al., 2016). In addition, the entire genome becomes more dynamic in response to DSBs, albeit to a lesser extent relative to repair sites (Krawczyk et al., 2012; Lottersberger et al., 2015; Mine-Hattab & Rothstein, 2012; Seeber, Dion, & Gasser, 2013). This global mobilization of the genome correlates to increased susceptibility to micrococcal nuclease digestion in response to DSB induction (Ziv et al., 2006), and possibly results from a global decrease in histone levels following DNA damage (Hauer et al., 2017; Kruhlak et al., 2006), release of chromatin-associated components (Ziv et al., 2006), or loss of anchoring to nuclear structures (Agmon, Liefshitz, Zimmer, Fabre, & Kupiec, 2013; V. Dion, Kalck, Seeber, Schleker, & Gasser, 2013; Strecker et al., 2016).

Understanding the molecular mechanisms involved in the relocalization of DSBs in heterochromatin and other DNA sequences requires the ability to analyze the 4D dynamics of repair sites. This can be accomplished by live imaging of repair components, given that many repair factors form cytologically visible foci upon recruitment to DSBs (Costes, Chiolo, Pluth, Barcellos-Hoff, & Jakob, 2010; Haaf, Golub, Reddy, Radding, & Ward, 1995; Lisby, Barlow, Burgess, & Rothstein, 2004; Lisby & Rothstein, 2004; Liu, Li, Lee, & Maizels, 1999; Maser, Monsen, Nelms, & Petrini, 1997; Ryu et al., 2015; Scully et al., 1997). For example, the repair component Mu2/Mdc1 associates with the phosphorylated form of the histone variant H2Av (Chiolo et al., 2011; Dronamraju & Mason, 2009; Stucki et al., 2005) (γH2Av, corresponding to mammalian γH2AX (Fernandez-Capetillo, Lee, Nussenzweig, & Nussenzweig, 2004; Madigan, Chotkowski, & Glaser, 2002)), a DSB mark, and mediates the recruitment of other HR proteins (Chapman & Jackson, 2008; Goldberg et al., 2003; Lou et al., 2006; Wang, Matsuoka, Carpenter, & Elledge, 2002). Thus, Mu2/Mdc1 foci can be used as a marker of repair sites throughout early and late steps of HR repair (Chiolo et al., 2011). ATRIP and TopBP1 are recruited to resected DSBs (Mordes, Glick, Zhao, & Cortez, 2008; Zou & Elledge, 2003), thus foci of these proteins mark resection. In late stages of HR, Rad51 promotes the search for a homologous template and strand invasion (Sung, 1994), and Rad54 stabilizes Rad51-mediated strand invasion intermediates (Petukhova, Stratton, & Sung, 1998; Petukhova, Van Komen, Vergano, Klein, & Sung, 1999). GFP-tagging of these components can thus be used as a marker for later repair steps (Figure 1C).

To investigate the spatial and temporal dynamics of heterochromatin repair, mEGFP-tagged repair components are monitored relative to mCherry (mCh)-tagged HP1a as a marker for the heterochromatin domain. Early studies revealed that Mu2/Mdc1, ATRIP and TopBP1 foci form inside the heterochromatin domain (Figure 1D, (Chiolo et al., 2011)), while Rad51 and Rad54 foci form after relocalization (Chiolo et al., 2011). Repair foci leave the domain primarily between 10 and 30 min after DSB induction with ionizing radiation (IR) (C. Caridi et al., 2017; Chiolo et al., 2011; Ryu et al., 2016; Ryu et al., 2015), and several foci reach the nuclear periphery in the first hour after IR (C. Caridi et al., 2017; Ryu et al., 2015). Thus, investigating the 4D dynamics of heterochromatic repair foci requires live cell imaging for at least 1 h after IR. A specific challenge of these studies is imaging the sample over long periods of time and with sufficient time points and Z-stacks to enable focus tracking, while limiting phototoxicity and photobleaching effects.

The use of *Drosophila* cultured cells greatly facilitates these experiments. In addition to the existence of a distinct heterochromatin domain, *Drosophila* cells are maintained at room temperature and ambient CO_2_ concentrations (Cherbas & Gong, 2014), which minimizes stress from environmental changes during cell culturing, sample processing, and live imaging. Further, these cells are mostly in S and G2 phases of the cell cycle (Chiolo et al., 2011), which is ideal for studying HR repair. Finally, efficient RNAi procedures facilitate the investigation of the mechanisms involved in heterochromatin repair with genetic approaches (Zhou, Mohr, Hannon, & Perrimon, 2013).

Here, we describe a procedure for monitoring the spatial and temporal dynamics of heterochromatic DSBs in *Drosophila* cells following IR (Figure 2), including: i) the generation of stable cell lines expressing fluorescent-tagged repair and heterochromatin marks; ii) how the same fields are imaged before and after IR; iii) the setup used to minimize light exposure with a DeltaVision deconvolution system (Applied Precision/GE Healthcare); and iv) post-image processing done with SoftWorX (Applied Precision/GE Healthcare), which maximizes the recovery of information from low-exposure experiments while correcting for modest photobleaching. Additionally, we describe the workflow we implemented to track repair foci with Imaris (Bitplane) and derive mean square displacement (MSD) curves with a Matlab (Mathworks) script. Finally, we describe a new method we developed in R to identify directed motions within mixed trajectories (*i.e.*, characterized by both diffusive and directed motions). Together, these techniques enable studying the spatial and temporal dynamics of heterochromatin repair, which cannot be accomplished with fixed cell studies or biochemical approaches. Similar approaches have also broad applicability for the study of nuclear dynamics of repair foci in other contexts, from yeast to mammalian cells.

**Figure 2:**
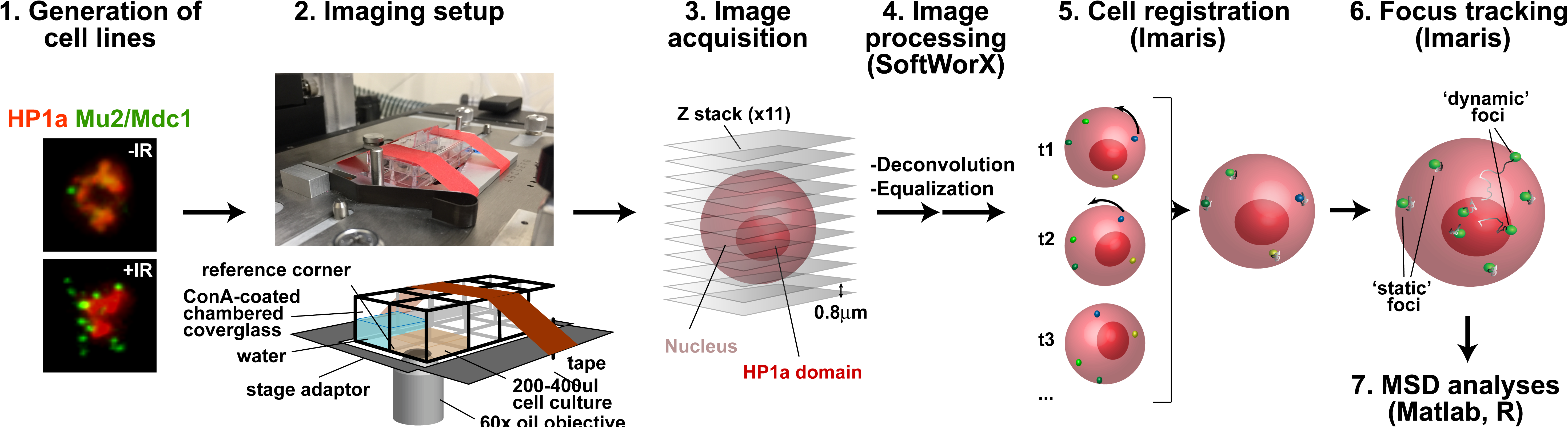
Pipeline for image acquisition, processing and analysis of focus dynamics in *Drosophila* heterochromatin. A stable cell line homogeneously expressing fluorescent tagged protein (*e.g.,* mCH-HP1a and mEGFP-Mu2/Mdc1), is placed in a well of a ConA-coated chambered coverslide, which is secured to the adaptor placed on the microscope stage. For long movies, evaporation of the media is limited by adding water to the surrounding wells. For tracking experiments, which require frequent imaging and long kinetics, photobleaching and phototoxicity are reduced by: i) selecting bright fluorescent tags, which reduce the exposure time required to detect repair foci and heterochromatin; ii) using the 2x2 binning option of the CoolsnapHQ2 camera, which dramatically increases the light collected by the camera per time unit; iii) underexposing the sample and recovering most image details with post-image processing. Individual nuclei are registered with Imaris to correct for modest rotational and translational shifts, using ‘static’ foci as a reference. ‘Dynamic’ foci are then tracked using Imaris and the biophysical properties of the motion are characterized using MSD analyses with Matlab and R.

## Methods

Successful focus tracking and 4D analyses of focus motion over long time periods requires the optimization of different steps, including the selection of the brightest and most photo-resistant fluorescent tags for live cell imaging, the use of microscopes that mitigate the risk of cell damage, and the identification of imaging conditions and post-imaging de-noising approaches to recover image details while minimizing light exposure. Cell mobilization and nucleus registration approaches are also needed to isolate focus dynamics from cell dynamics. Once the positional data are collected, quantitative analyses enable the understanding of the biophysical properties of focus motion, including identifying directed motions.

### 1. Live Cell Imaging of Drosophila cells

#### 1.1 Generation of stable cell lines expressing fluorescent-tagged proteins

Live cell imaging experiments benefit from using stably transfected Kc167 (Kc) cells maintained as exponentially growing cultures. Kc cells are preferred over S2 cells because they adhere better to the substrate and maintain a more stable karyotype in the population. Live imaging and tracking experiments are facilitated by generating cell lines with homogeneous signals across the cell population, and a wise selection of tags for live imaging.

##### 1.1.1. Cell maintenance

Kc cells are maintained in Schneider’s medium (Sigma) supplemented with 10% FBS (Gemini) and 2% Antibiotic-Antimycotic (Gibco) at 27°C. Schneider’s media is an optimal choice for live imaging given the low autofluorescence. If cells are grown in media with high autofluorescence (e.g., SF-900 II, Gibco), we recommended shifting the cells to Schneider’s media just before live imaging. Cells are kept in exponential phase by splitting the culture every 3-5 days to maintain a concentration of 1.5-9 x 10^6^ cells/ml. Cells do not grow equally well when seeded at a density below 1.5 x 10^5^ cells/ml. For information about *Drosophila* cell maintenance see the *Drosophila* Genomics Resource Center (DGRC) website and (Yang & Reth, 2012).

#### 1.1.2 Selecting fluorescent tags for live cell imaging

Live cell imaging experiments are critically dependent on selecting the best combination of fluorescent tags for each protein of interest to maximize signal recovery during the entire kinetic. We have successfully used several fluorescent tags for live imaging of *Drosophila* cells (*i.e.,* EGFP, GFP, mEGFP, mCitrin, mCerulean, mTurquoise2, mCherry, Aquamarine). However, mEGFP or EGFP tags signals are among the brightest and most resistant to photobleaching, and have been the best choice for 4D tracking of repair foci that requires frequent imaging over long periods of time (Ryu et al., 2015). mCherry (mCh) is also quite resistant to photobleaching and can be used to detect very abundant proteins or large nuclear structures in the same experiments, such as the heterochromatin domain (*e.g., via* expression of mCh-HP1a) or nuclear periphery components (*e.g., via* expression of mCh-LaminC) (Chiolo et al., 2011; Ryu et al., 2016; Ryu et al., 2015).

##### 1.1.2 Generating cell lines that express tagged proteins

Generating *Drosophila* cell lines that express fluorescent markers for DSB repair and heterochromatin is facilitated by using agents that deliver high transfection efficiency, like Cellfectin (Invitrogen), Transit-Insect (Mirus Bio LLC), Transit-2020 (Mirus Bio LLC), or Effectene (Quiagen) (Figure 3). Transfections are done following manufacturer's procedures in 6-well plates and typically using 2.5 µg of plasmids expressing each tagged protein of interest. Transfections with up to three plasmids result in most cells expressing all plasmids. For live imaging experiments requiring transient transfection, Cellfectin is preferred because it forms few if any precipitates during transfection. Transgene expression can be tested 3-4 days after transfection.

**Figure 3:**
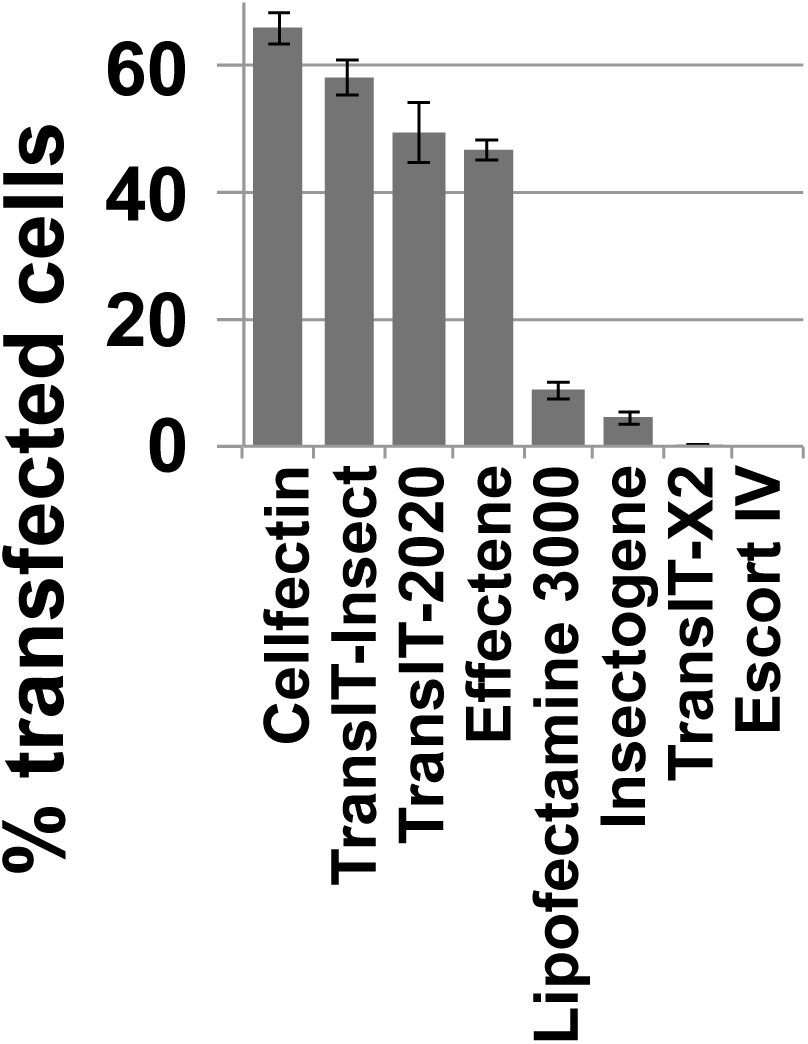
High transfection efficiency is obtained with Cellfectin. High transfection efficiency facilitates the generation of stable Kc cell lines expressing fluorescent markers of the heterochromatin domain and damage foci. To optimize this step, different transfection agents were tested following manufacturers’ instructions. Cells were transfected with 2.5 µg of a midiprep of pCopia-mCh-HP1a and the average number of cells expressing HP1a was estimated 72 h after transfection. The highest transfection efficiency obtained after several optimization steps is shown for each reagent, revealing the highest transfection efficiency using Cellfectin. Error bars show +/- SEM.

Using stable cell lines facilitates acquiring significantly more data in a short time, as every imaged field will contain ~20-50 transfected cells that can be analyzed for focus dynamics. Additionally, some repair components and nuclear architecture markers are not detectable after transient transfection, in which case stable cell lines are required. This needs to be empirically determined. To generate stable cell lines, 1 µg of plasmid carrying the selection cassette (*i.e.,* pCoPuro (Iwaki, Figuera, Ploplis, & Castellino, 2003) or pCoHygro (Invitrogen)) is added to the plasmid mix during transfection, and cells are split in media containing the selection agent (*i.e.,* 150 µg/ml Hygromycin or 300 µg/ml Puromycin) 3-5 days after transfection. For the following weeks, cells are split as needed to maintain them in exponential growth phase, and recovery from the selection is typically observed after 4-5 weeks. Longer selection times render the signal more homogeneous across the culture but might also result in reduced signal intensity. Thus, for tracking experiments we recommend using the cells shortly after establishing a stable line.

Notes: We mostly use the Copia promoter to constitutively induce a protein of interest at low levels (typically similar to endogenous levels), although the metallothionein promoter (MT, induced by CuSO4) and the heat-shock promoter (hs, induced by a 30 minute temperature shift at 37°C) have also been used successfully. Given the potential effects of temperature shifts on gene expression, nuclear architecture, chromatin dynamics, or repair pathways (Li et al., 2015; Seong, Li, Shimizu, Nakamura, & Ishii, 2011; Velichko, Petrova, Kantidze, & Razin, 2012), and the potential secondary effects of expressing non-physiological levels of the proteins, specific controls need to be applied when using these promoters. Additional considerations apply to multi-color imaging. Imaging more than two channels increases the risk of phototoxicity and the time required for imaging each field. These might particularly affect tracking experiments, which require many time points and frequent imaging. Additionally, transfections with four plasmids (three plasmids expressing the tagged proteins plus the plasmid carrying a selection marker) are not very efficient, which complicates the generation of stable cell lines expressing all three fluorescent markers. If three color imaging is required, it is recommended to express two (or three) of the tagged proteins in a multi-cistronic vector (Gonzalez et al., 2011). Alternatively, the selection marker can be cloned into one of the three vectors, or multiple tagged components can be integrated in the genome with sequential transfections.

#### 1.2. Immobilization of cells for microscopy

A key step for 4D analysis of nuclear dynamics is immobilizing the cells on the substrate. This minimizes rotational and translational movements of the cells, reducing the need for post-imaging corrections of the movement and dramatically improving the analysis of nuclear dynamics. We tested ConcanavalinA (ConA) (Type VI or IV, Sigma), Polylisine (Sigma), and Cell-Tak (Corning), and found that ConA Type VI produces the best results.

Procedure for coverslide coating with ConA:

1.2.1 Prepare a solution of 1 mg/ml ConA in water. Stir for about 1 h, until mostly dissolved. Filter the solution with a 0.22 µm pore-size filter.

1.2.2 Add 100 µl of ConA solution to each well of the 8-well chambered imaging coverslide (Nunc™ Lab-Tek™ II Chambered Coverglass, Thermo Scientific). Let the solution dry in the hood or in a 30°C incubator. For best results repeat the coating 2 more times and use the coated coverslides within one week of preparation.

#### 1.3. Temperature regulation during live imaging

Maintaining a stable temperature while imaging is important to ensure consistency throughout the time course as well as across different experiments. To achieve stable temperature conditions, we use an environmental chamber mounted around the microscope that maintains the sample at 25°C during the experiment.

Note: For movies lasting a few hours, it is also important to limit evaporation of the growth media by adding water to empty wells surrounding the well with the cells (Figure 2). For even longer movies, it is helpful to place a slightly damp paper towel in the empty chambers to further limit evaporation.

#### 1.4. Image acquisition setup with a DeltaVision deconvolution microscope

Depending on the proteins being analyzed and the microscope available, imaging parameters need to be empirically optimized to minimize photobleaching and phototoxicity. For imaging the dynamics of heterochromatic repair sites, we typically utilize cells stably expressing mCh-HP1a and a repair protein tagged with mEGFP (e.g., Mu2/Mdc1, ATRIP or Rad51 (Chiolo et al., 2011; Ryu et al., 2016; Ryu et al., 2015)). The following procedure is optimized for cells expressing mCh-HP1a and mEGFP-Mu2/Mdc1 and imaging with a DeltaVision Elite deconvolution inverted microscope equipped with: white light LED (rated 100 Lumens at 350 mA); 7-color InsightSSI solid state illumination system; PlanApo 60x oil objective with N.A. 1.42; Ultimate Focus module; a Coolsnap HQ2 camera; and controlled by SoftWorX software (v. 6.1.3). The Deltavision System is optimized for low-light imaging and image processing through deconvolution, which enables an excellent recovery of image details in underexposed samples (see 2.1). This makes this system an excellent choice for the experiments described here. Spinning disk microscopes or widefield microscopes used in combination with external image processing software have also been used successfully for similar experiments (see (Hediger, Taddei, Neumann, & Gasser, 2004; Meister, Gehlen, Varela, Kalck, & Gasser, 2010) for an overview of different imaging technologies and (Cho et al., 2014; Dimitrova, Chen, Spector, & de Lange, 2008; Vincent Dion et al., 2012; Lottersberger et al., 2015; Mine-Hattab & Rothstein, 2012; Su et al., 2015) for examples of their application to focus tracking experiments).

1.4.1 Split cells to a density of 2 x 10^6^ cells/ml 48 h before the experiment.

1.4.2 On the day of the experiment, transfer 200-400 µl of cells into one well of the chambered coverslide, and let the cells settle for 10-15 min before imaging. Meanwhile, set the temperature of the environmental chamber of the microscope to 25°C.

1.4.3 On the microscope, manually position the 60x objective and the dichroic filter for GFP/mCh imaging.

1.4.4 Place immersion oil on the objective lens. Note that the immersion oil needs to be optimized based on objective, temperature conditions and coverslip thickness to minimize spherical aberration while maximizing contrast in the images. With 1.5 mm chambered coverslide we use an immersion oil refractive index=1.512 (GE Healthcare Life Sciences). Follow the microscope manufacturer’s instructions for this step.

1.4.5 Place the chambered coverslide on the stage adaptor. When imaging multiple fields, the movement of the stage can shift the coverglass on the stage adaptor, making it difficult to return to the same field of cells. To prevent this issue, secure the chambered coverslide tightly using tape (Figure 2) before placing it on the microscope stage. Adjust the stage level to bring the sample into focus.

1.4.6 In an effort to minimize light exposure during imaging, thus reducing photobleaching and phototoxicity, set the Coolsnap HQ2 camera at 2x2 binning. This results in less resolution but higher intensity collected per pixel. With this setting, we set an image size of 512 x 512 px, to collect the largest possible field of view.

1.4.7 Select the fields of interest. Illumination intensity is adjusted to the minimum level sufficient to see the sample to minimize photo-bleaching of the signal while choosing the fields (*e.g.,* 10% excitation intensity, or 10%T, for mCh and GFP). Suitable fields should contain an even distribution of cells as a mono-layer, in addition to intense and homogenous signals for mCh and GFP-tagged proteins. The number of fields that can be imaged in a single experiment is limited by the time required for imaging each field and the time interval between time points. We typically image 4-5 fields for each experiment. Once each field is selected, save its coordinates using ‘*Mark Point’* option in the ‘*Point List’* section in SoftWorx. The list of selected fields will appear in the '*Point List'* window.

1.4.8 Optimize the imaging path to visit all the fields using ‘*Optimize List’* followed by ‘*Compact List’* commands in the ‘*Point List’* section. This will minimize the time required to visit all of the selected fields, enabling imaging of more fields for each experiment.

1.4.9 Because cells are imaged before and after IR, it is essential to be able to return to the same fields after removing the chambered coverslide from the microscope for IR exposure. However, removal and repositioning of the sample might result in a slight shift of the stage (Figure 4). To identify the same field of cells, select a ‘landmark’ field to use as a reference (such as a corner of the slide), and save its coordinates in addition to those of the selected fields of cells (Figure 4).

**Figure 4:**
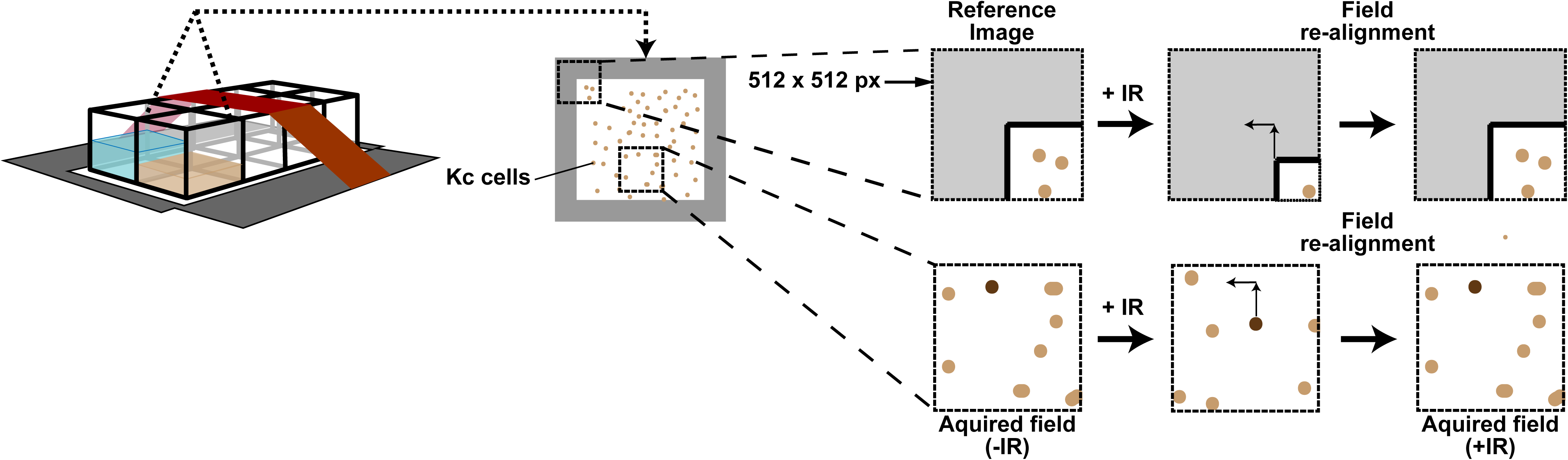
Field realignment post IR. Focus tracking requires cell imaging before and after IR, but repositioning of the chambered coverslide on the stage after IR frequently results in a slight shift of the stage. This can be corrected using a reference field (*e.g.,* a corner of the well). The reference field is manually re-centered after IR, and the extent of correction applied enables determination of the X and Y shift. Corresponding corrections are applied to all fields of interest.

1.4.10 Select the time of exposure for the sample based on the need to detect sufficient signal with the minimum exposure. *E.g.,* we use 10% T and a target intensity of 200 and 600 counts (approximately 5 and 15 msec) for mCh-HP1a and mEGFP-Mu2/Mdc1, respectively. This results in underexposing the image, but most image details can be recovered post-imaging by optimized deconvolution and photobleaching correction algorithms available in SoftWorX (Figure 5 and Section 1.5 of this protocol).

**Figure 5:**
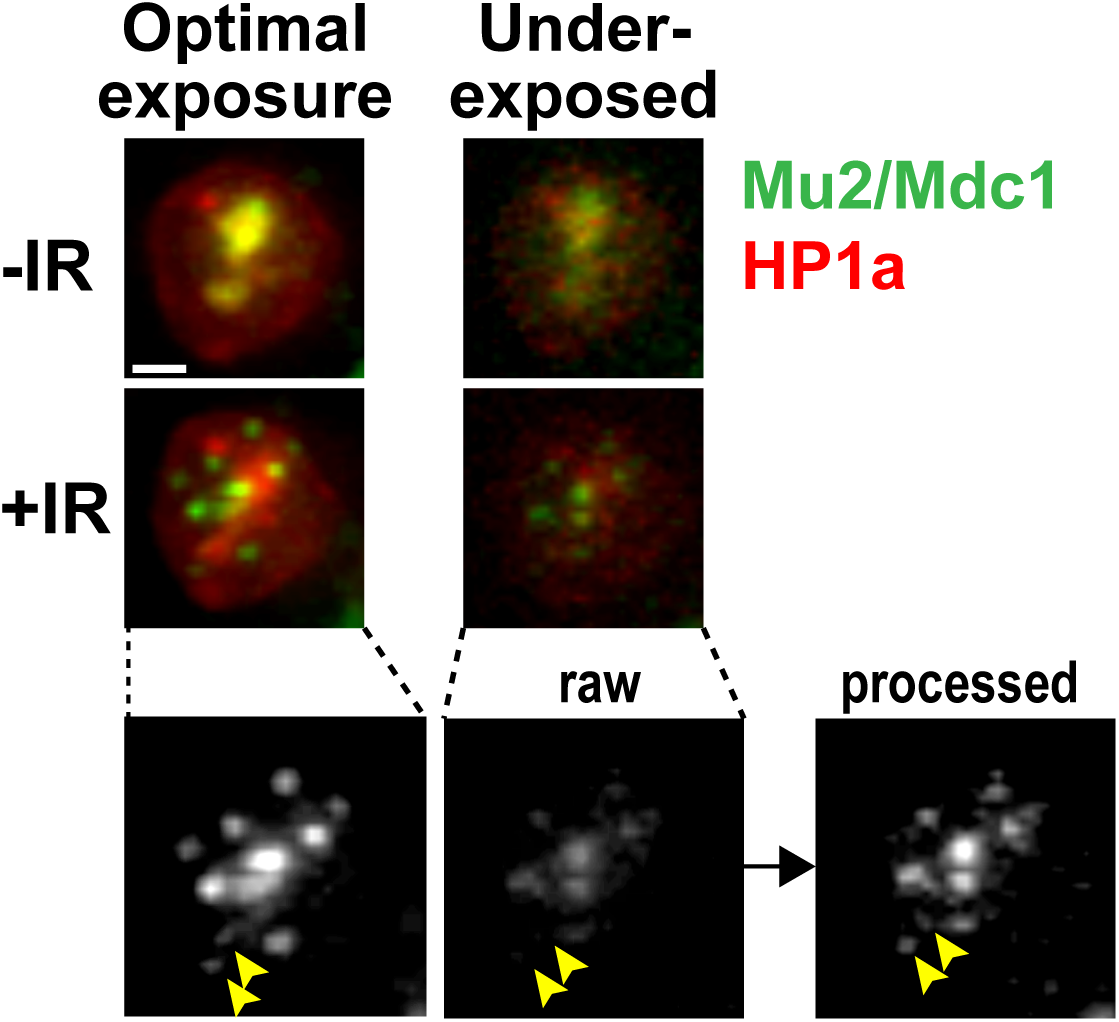
Image details in underexposed samples are recovered with post-image processing. Underexposure of the cells enables the long and frequent image collection required for focus tracking while minimizing cell damage and photobleaching. The application of deconvolution and equalization algorithms enables recovery of most details in underexposed images. In the example shown, a Kc cell stably expressing mEGFP-Mu2 and mCh-HP1a shows an overall enrichment of Mu2/Mdc1 signals inside the heterochromatin domain before IR (-IR), and damage foci associated with the heterochromatin domain at 10 min after IR (+IR) (Ryu et al., 2015). The signal has been collected first in underexposed conditions (5 ms for mCh and 20 ms for GFP, with 10% T), then in optimal imaging conditions (12 ms for mCh and 20 ms for GFP, with 100% T), for both time points as indicated. The noisy signal of raw underexposed images is significantly improved by post-image processing (deconvolution and equalization), revealing even the weak signals associated with small foci (arrowheads). Images are max intensity projections of one nucleus. Scale bar = 1µm.

1.4.11 Adjust image acquisition settings to image the sample across its entire thickness with a minimum number of Z stacks. This requires some optimization depending on the thickness of the sample. For *Drosophila* cells, image 11 Z-stacks at 0.8 µm distance between the Z-stacks. While a relatively high distance between Z stacks reduces the resolution in this dimension, this is an excellent compromise between spatial resolution required for tracking and the need to minimize photobleaching and phototoxicity.

1.4.12 List the fields of interest (*i.e.,* the points saved in the '*Point List'* section) in the ‘*Design/Run Experiment’, ‘Points’* tab, ‘*Visit Point List’* section. Activate the ‘*Ultimate Focus’* options with 5 iterations to assure that cell focus is maintained throughout the experiment. This option corrects for axial drifts with an infrared laser-based system that detects the position of the coverslip relative to the sample. This is preferred over the ‘*Image-based autofocus before imaging’* option which causes photo-damage while refocusing the image. If the ‘*Unlimited Focus’* module is not available on the system, manual adjustments of the focus or automated image-based autofocusing might be required throughout the kinetic.

1.4.13 Begin image capturing using ‘*Run’*-‘*Start Scan’* (green arrowhead in the ‘*Design/Run Experiment’* window). The acquired image set corresponds to the ‘untreated’ time point (-IR, Figures 1-4).

1.4.14 Once all frames have been imaged, carefully remove the chambered coverslide without disturbing the stage position or the cells. Apply additional immersion oil onto the objective lens as needed.

#### 1.5. IR treatment, field re-alignment, and image acquisition after IR

This procedure enables the acquisition of several selected fields of cells after IR for each experiment, providing a large number of cells for analyses. Importantly, a direct investigation of cellular responses to IR requires the ability to image the same fields before and after IR, which is accomplished as described below.

1.5.1 We routinely expose the sample to 1.7 or 5 Gy IR, which corresponds to 14.5 s or 44 s in our x-ray irradiator (X-RAD iR-160, Precision X-Ray, stage level 30), respectively. The timer is started half way through IR exposure. We use 5 Gy to track ATRIP foci (Ryu et al., 2015), and 1.7 Gy for Mu2/Mdc1 foci. Mu2/Mdc1 foci are typically more numerous than ATRIP foci, so reducing the intensity of IR minimizes overlapping signals (*e.g.*, deriving from focus clustering (Chiolo et al., 2011; Chiolo et al., 2013), reducing the risk of ambiguous tracks. Notably, we did not detect major differences in MSD values between 1.7 and 5 Gy in our cells.

1.5.2 After exposure to IR, carefully place the chambered coverslide back on the microscope stage. Position the sample on the reference field (*e.g.,* the corner of the chambered coverslide) using its saved coordinates (Figure 4). Compare to the untreated images. If the stage shifted, move back the reference field until the original position and record the extent of the shift required for this realignment. Readjust the position of each field of cells accordingly. For example, if the reference field is shifted +300 µm along the X axis and +400 µm along the Y axis, move each field of cells -300 µm and -400 µm along the X and Y axes, respectively, and save the new position (Figure 4). For each field, also readjust the Z axis to ensure that the sample is in focus and save the new position.

1.5.3 Adjust the ‘*Image Capturing’* parameters of the ‘*Experiment Setup’* option to collect images at 40 s time intervals for 1 h movies (91 images total). In SoftWorX, these options are under the ‘*Time-lapse’* tab in the ‘*Design/Run Experiment’* window. Click ‘*Start Scan’* to begin the imaging of the time points after IR (+IR, Figure 2). Record the time point from IR at which the imaging starts (this is 3-5 min after IR in our experiments). Let the system acquire all the time points.

Notes: The imaging protocol described has been optimized for live imaging and tracking for MSD analysis of focus motion in response to IR. However, similar imaging approaches have broader applicability. For example, damage can be induced with hydroxyurea (1-2 mM) or methyl methanesulfonate (0.0033-0.1%). If cells are treated with chemicals, it is recommended to image untreated cells under control conditions to check for cellular toxicity coming from the solvent or imaging procedures. Longer movies are typically collected with longer time intervals between images to minimize photobleaching and phototoxicity. For example, we acquire images every 80 s for 2 h movies and every 30-60 minutes for 24 h movies. Finally, it is necessary to optimize imaging parameters (number of Z-stacks, intensity of light during exposure, frequency of time points) for each cell line, tagged component, and treatment. We recommend testing the effect of the imaging approach *per se* on DSB formation (e.g., Mu2/Mdc1 focus number (C. Caridi et al., 2017)) and cell ability to divide (*e.g.,* by following the cell overnight and capturing cell division by imaging with 40 min time intervals (Chiolo et al., 2011)). Non-invasive imaging approaches should not interfere with cell division or induce DSB formation (Meister et al., 2010).

### 2. Image processing and focus tracking

#### 2.1 Image processing with SoftWorX

The low light exposure conditions used in this protocol enable imaging over long time periods while limiting phototoxicity and photobleaching, but they also result in low signal to noise ratio in the collected images. A critical step in this analysis is the application of algorithms (deconvolution and equalization) that recover most image details post-imaging, enabling precise identification and tracking of repair foci throughout the entire kinetic. Equalization compensates for photobleaching effects by normalizing the signal intensity of each time point to a reference time point. This greatly facilitates automated detection and tracking of foci, which relies on the average signal intensity across the kinetic. Deconvolution is the mathematical correction of the distortion of an image resulting from diffraction and aberration of light passing through the optics of a microscope (Scalettar, Swedlow, Sedat, & Agard, 1996). This distortion can be quantified by establishing the shape of a single point object imaged with a microscope, or point spread function (PSF) (Scalettar et al., 1996). Deconvolution algorithms use the PSF to correct for the distortions, resulting in image deblurring, noise reduction, resolution improvement and contrast enhancement (Scalettar et al., 1996). In addition to facilitating automated focus detection by increasing resolution, deconvolution corrects for image distortions in the axial direction, dramatically reducing noise in the tracks coming from this dimension. In the DeltaVision microscope, equalization and deconvolution algorithms are fully integrated with the system and optimized for the specific microscope setup. However, similar results can be obtained with other software (*e.g.,* Huygens Software Suite for deconvolution or Imaris for equalization).

2.1.1 Combine “-IR” and “+IR” files using the ‘*Image Fusion’* function in SoftWorX with the “C*ombine time points for like wavelengths”* option.

2.1.2 Deconvolve the fused images in SoftWorX using 5 iterations of the ‘*Conservative’* protocol to improve contrast and resolution. In our experience this is sufficient to improve image sharpness while limiting potential artifacts.

2.1.3 Correct the deconvolved file for modest photobleaching using the '*Equalize Time Points'* option.

2.1.4 Having individual cells cropped facilitates further image analysis with Imaris. For this purpose, crop selected cells using the '*Save File'* option in SoftWorX by selecting the field section of interest. Cells better suited for the registration step described next need to fit the following criteria: i) the cell should appear nearly static throughout the kinetic; ii) at least 4 ‘*static’* foci (*i.e.,* foci that don’t significantly move in the nucleus) are present throughout the kinetic (Figure 2) - these will be used for registration; iii) GFP and mCherry signals remain visible throughout the kinetics.

#### 2.2 Cell registration with Imaris

After cropping the selected cells, we utilize Imaris (v. 7.7.1 with XT module) to correct for minor cell/ nucleus movement (‘registration’) and to track foci for motion analysis. Registration is performed by tracking all of the foci (usually 4-12) that remain largely static throughout the kinetic and by correcting cell drift using those as a reference (Figure 2). Using 7+ foci as a reference typically results in a better registration.

##### 2.2.1 File cropping

Remove the first time point (UNT) at this stage of the analysis by using the '*Crop Time'* function in Imaris. The first time point does not contain any IR-induced focus, so it cannot be used for registration. Save the cropped file with a new name to use this for further analysis. Keeping the original file is also important, as it contains information about which repair foci were already present before IR. Those foci can be used for registration but not for tracking of IR-induced foci and MSD analyses.

##### 2.2.2 Optional: noise reduction

Automatic focus tracking is greatly improved by reducing background signals and small focus vibrations along the Z stack. Background signals are reduced using Imaris, by applying the '*Baseline Subtraction'* option in the '*Thresholding'* function of '*Image Processing'* to the channel with the foci. In addition, minor vibrations along the Z-stack are reduced by applying the '*Smooth Time'* function of '*Image Processing'* with a filter width of 1 to all time points. Together, these steps facilitate the subsequent automated tracking of foci.

##### 2.2.3 Automated focus tracking

Focus tracking is done using the Imaris ‘*Spot Detection Tool’*, and the tracked foci are used to register the nucleus. Apply the following steps to generate the tracks, clicking the right pointing blue arrow to proceed through each step:

i. Generate a new ‘Spot’ in Imaris. Select the magic wand icon, and click on ‘*Rebuild’*;
ii. Select the “*Track Spots Over Time”* box;
iii. Select the ‘*Source Channel’* corresponding to the wavelength at which foci were imaged, using the dropdown menu;
iv. In the ‘*Spot Detection’* section, select 0.2 µm as ‘*Estimated XY Diameter’*. This value reliably detects most DNA repair foci we examined. However, depending on the proteins being studied, this parameter may require empirical adjustments;
v. The algorithm will place spheres corresponding to all detected foci. In the ‘*Filters’* section select ‘*Quality’*. Adjust the lowest threshold to a point at which the faintest foci are reliably distinguished from the background;
vi. In the ‘*Add/Delete (Cursor Intersects with)’* section, using the dropdown menu, select the ‘*Specific Channel’* corresponding to the wavelength at which foci were imaged;
vii. In the “*Algorithm”* section, select the ‘*Autoregressive Motion’*. In the “*Parameters”* section change the ‘M*ax Distance’* to 0.5 µm and the ‘*Max Gap Size’* to 3. Check the box labeled ‘Fill gaps with all detected objects’;
viii. We apply two filters in the “*Classify Tracks”* section. Select ‘*Track Duration’* and adjust the lower threshold to eliminate tracks that only last a few time points. Add the ‘*Track Length'* filter and adjust the lower threshold to further remove short tracks. Click the right pointing orange double arrow icon to finalize the track detection.

##### 2.2.4 Manual editing of tracks

Sometimes the tracks generated by Imaris include large jumps to unrelated foci, especially when those are in close spatial proximity relative to the focus of interest, in which case tracks detected automatically require manual adjustments. To edit a track, select the corresponding spot and the‘*Edit Tracks’* icon. Select the time point that requires editing, delete it, manually recreate a new spot and connect it to the pre-existent track. Edit each track as necessary to assure that each focus is correctly identified throughout the kinetic.

##### 2.2.5 Selection of tacks for registration

Identify foci that remain relatively stationary throughout the time course. These are typically euchromatic foci that originate outside the HP1a domain or foci that were already present in the nuclei before IR (*e.g,* spontaneous damage). We find that registration works better using at least 7 foci, although 4 foci are sometimes sufficient. Foci characterized by extensive motion will affect the registration process and should be excluded at this stage.

##### 2.2.6 Registration

Highlight all suitable tracks and click the ‘*Correct Drift’* button below the tracks window. In the ‘*Drift Correction Options’*, select ‘*Translational And Rotational Drift’*. For the ‘*Result Dataset Size’* select ‘*Include Entire Result’*. Confirm that the “*Correct objects’ position*s" box is selected. Then click ‘OK’. Imaris will register the nucleus based on the selected tracks, which will compensate for any minor translational and rotational motion of the nucleus during the experiment. Save the resulting file with a new name. This will be used for further focus tracking (Figure 2).

##### 2.3 Focus tracking with Imaris

Tracking DNA damage foci is similar to the registration process, except that a new ‘spot’ is generated for each tracked focus.

##### 2.3.1 Optional

To minimize artificial vibrations of the nucleus resulting from the registration process, reapply the '*Smooth time'* function of '*Image Processing'* with a filter width of 1 to all the time points.

##### 2.3.2 Focus tracking

Visually identify a focus for tracking. Repeat steps 2.2.3-2.2.4, except that most tracks are filtered out using both upper and lower thresholds for '*Quality'* and '*Track Duration'* filters; only the focus of interest remains tracked. Repeat this step as many times as necessary to track all foci under investigation.

##### 2.3.3 Nuclear periphery detection

The nuclear periphery is identified by creating a volume that corresponds to the diffuse nuclear signal generated by background mCh-HP1a or GFP-Mu2/Mdc1 signals. Using the ‘*Automatic Creation’* function, select the channel corresponding to HP1a or Mu2/Mdc1, and manually adjust smoothness and threshold to create a volume fitting the nuclear signal. Alternatively, a specific marker for the nuclear periphery (*e.g.*, mCh-LaminC) can be used (Ryu et al., 2015).

#### 2.4 4D image rendering

##### 2.4.1 Optional

4D rendering of individual tracks can be done in Imaris to facilitate the analysis and display of each track (i.e., Figure 1D). 4D rendering is obtained by generating a volume corresponding to the HP1a domain. Using the ‘*Automatic Creation’* function, select the channel corresponding to the HP1a domain, manually adjust smoothness and threshold to create a volume fitting the HP1a signal. The volume corresponding to the nuclear periphery (defined as described in 2.3.3) can be displayed at this step as well. Finally, select the focus track of interest, and deselect the green and red channels of the original image before saving the image.

### 3. Analysis of focus dynamics

#### 3.1 MSD analysis

MSD analyses, which plot the average squared distance traveled by a focus at progressively increasing time intervals, provide quantitative measurements of the dynamic properties of focus motion (V. Dion & Gasser, 2013; Saxton & Jacobson, 1997; Spichal & Fabre, 2017). MSD values are calculated at multiple time intervals to generate a curve for each track, and these curves are averaged to describe the behavior of a population of foci. The shape of the MSD curve reveals whether the motion of a particle is Brownian, sub-diffusive or directed (V. Dion & Gasser, 2013; Saxton & Jacobson, 1997; Spichal & Fabre, 2017). MSD curves with increasing slope reflect directed motion, while linear MSD graphs indicate Brownian motion (V. Dion & Gasser, 2013; Meister et al., 2010; Saxton & Jacobson, 1997; Spichal & Fabre, 2017) (Figure 6A). However, given that the chromatin behaves as a polymer, and other constraints to the movement exist (*e.g.,* chromatin compaction, molecular crowding, and anchoring to nuclear structures (V. Dion & Gasser, 2013; Spichal & Fabre, 2017)), chromatin motion is typically sub-diffusive rather than Brownian, resulting in flattened MSD curves (Spichal & Fabre, 2017) (Figure 6A). In addition, sub-diffusive (or Brownian) motion occurring in a constrained space (*e.g.,* the nucleus or subnuclear domains) are characterized by MSD graphs that reach a plateau (V. Dion & Gasser, 2013; Saxton & Jacobson, 1997; Spichal & Fabre, 2017), and this is proportional to the radius of constraint (*i.e.,* the radius of the volume explored by the focus) (Figure 6B). MSD curves enable calculating the radius of constraint and the diffusion coefficient as follows.

**Figure 6:**
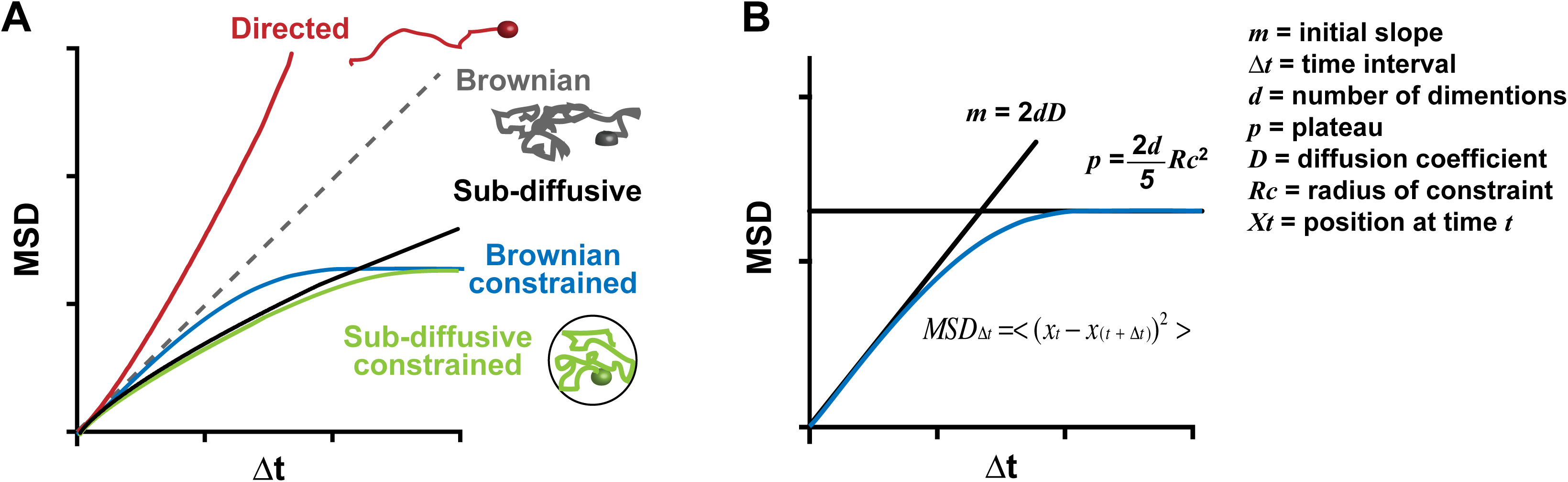
MSD curves. MSD values are derived from the mean of the squared displacement calculated over increasing time intervals, Δ*t*. A) Examples of MSD plots corresponding to directed motion, Brownian motion, sub-diffusive motion and sub-diffusive or Brownian motions via a constrained space are indicated. B) For Brownian or sub-diffusive motions, containment radius and diffusion coefficient are calculated using the plateau of the curve or the initial slope, as indicated (see text for details).

##### 3.1.1 Extracting positional data

For each tracked spot, select the ‘*Statistics’* tab and click on the ‘*Position’* information in the dropdown menu of the ‘*Detailed’* tab. Click on the floppy disk icon to save the data as an excel file. The file contains three columns: posX, posY, posZ, corresponding to the coordinates of the focus at each time point. To use the MSD script provided as Supplementary Information 1: ‘MSD_script.’, add a column before those three and name it ‘t’. Number each time point with increasing number starting from ‘001’ for the first time point, add the corresponding number at the beginning of each file name, and save this as a comma-separated values comma-separated values (.csv) file editable in Excel.

##### 3.1.2 MSD calculation

To derive MSD curves, positional data obtained from each track are processed with a Matlab script that generates MSD values for each focus. Open the MSD script in Matlab and point to the folder containing the .csv files in line 2 (*e.g., ‘myInputFolder’*). Enter the information of the destination folder in line 3 (*e.g., ‘myOutputFolder’*). Adjust the number of time points to be analyzed by editing the *timelapse_length* variable in line 8 (e.g., 92 timepoints in our experiments). This change will be reflected in the MSD table size and calculation. MSDs were calculated as described in (V. Dion & Gasser, 2013; Meister et al., 2010; Miné-Hattab & Rothstein, 2012; Spichal & Fabre, 2017); for each position ***x***_*t*_ (characterized by XYZ coordinates) at time ***t***, and time interval ∆***t***:

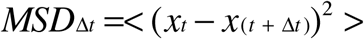

Run the script. Output files will be saved in the destination folder. Note that multiple files (*i.e.,* positional data from a large number of foci) can be read at once. As long as the file name is preceded by a progressively increasing number (*i.e.* 001, 002, etc)., the output will be a .csv file, with columns corresponding to MSD values for each focus and the focus number indicated at the bottom of each column. Rows correspond to increasing ∆***t*** (i.e., ∆***t***=1 in row 1, ∆***t***=2 in row 2, etc.). Combine all MSD data in one Excel file. Calculate and plot the average MSD value and standard errors.

##### 3.1.3 Calculation of the radius of constraint and the diffusion coefficient for a population of foci

In order to calculate the plateau of the MSD curve, derive the curve that fits the data using Matlab’s ‘*curve fitting tools’* with this equation:

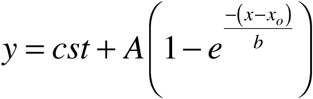

The plateau ***p*** of the curve, corresponding to (***p*** = *cst +A),* enables to calculate the radius of constraint ***Rc***
as:

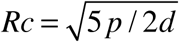

where ***d*** is the number of dimensions (3 in this case). The diffusion coefficient ***D*** is derived from:

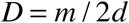

where ***m*** is the initial slope of the MSD curve.

#### 3.2 Detection of directed motions (DM) in mixed trajectories

When directed motion occurs in a trajectory characterized by different types motion (*e.g.,* preceded and/ or followed by sub-diffusive or constrained motions), also called mixed trajectory, an MSD analysis applied to the entire kinetic might mask the presence of directed motions. This is even more likely when MSD values across different foci are averaged, using directed motions initiated at different time points for each focus. Here we describe a method to identify time intervals characterized by directed motions in these mixed trajectories, for each focus track. MSD curves for individual tracks are analyzed through a script in R (Supplementary Information 2: ‘DM_script.R’) that returns the time intervals characterized by MSD graphs with increasing slope.

##### 3.2.1 DM identification

Open the script in R (cran.r-project.org) or R studio (rstudio.com) and point to the folder containing one of the .csv files obtained in section 3.1.1, in line 202 (*i.e., ‘list.files’*). The script derives MSD curves starting from each time point of the trajectory, and with progressively increasing ∆***t*** (with ∆***t*** >10 time intervals). MSD curves are then smoothed using the *lowess* function with default parameters (stat.ethz.ch/R-manual/R-devel/library/stats/html/lowess.html) and segmented in two or more lines with increasing slopes. Points of slope change along the curve are detected using a *change point* analysis (Zeileis, Kleiber, Krämer, & Hornik, 2003) (*breakpoints* function with parameter fraction=0.3; *i.e.* every regression fit must use at least 30% of the total data points). The ratio of slopes between two lines is used to identify directed motion, *i.e.* time intervals when the MSD graph shows progressively increasing slopes. A few parameters can be modified to optimize the detection of directed motions; the current values robustly detect directed motions in our datasets but adjustments might be required for different data:

i. *‘minratio’* is the minimum ratio between lines derived from MSD plots for considering the graph characterized by increasing slope, and the current value is set at 3.5. Higher values of *minratio* increase the stringency of detection of directed motions (Note: a *minratio* higher than 2 is required to minimize the detection of false positives).
ii. *'r2min’* is the minimum coefficient determination of the fitted line. This is set at 0.99 and higher values of this parameter increase the stringency of detection of directed motions.
iii. *‘minslope’* is the minimum slope of the regression line at which data points are considered (current setting: 0.012). Lower values result in more foci with limited motions included in the analysis.

Run the script with the selected parameters. Time intervals characterized by directed motion appear in the ‘*Console’* work space and the corresponding graph is shown in the ‘*Plots’* section. If parameters are set correctly, those graphs show an upward curvature indicating directed motions. The first and last time point of each time interval characterized by directed motion are saved in a file generated with file extension ‘.out’ defined in the ‘*outsuffix'* variable, and are shown as time intervals. Data are compiled to show the number and duration of directed motions across the kinetics. Those typically correspond to time intervals characterized by directed motion in a mixed trajectory. The detection of time intervals characterized by directed motion can be validated by running the MSD script described in section 3.1 on the positional data corresponding to each time interval.

## Acknowledgments

This work was supported by R21ES021541, the Mallinckrodt Foundation Award, American Cancer Society Research Scholar Grant, and R01GM117376 to I.C. We would like to thank S. Keagy for insightful comments on the manuscript, B. Spatola for his help on an early version of the manuscript, T. Nellimoottil for his help with the MSD script, T. Ryu for her analysis of Imaris filtering options and ConA optimization experiments, K. Chrzanowski for his input on Imaris tools, and the Chiolo Lab for helpful discussions. We also thank R. Rothstein, J. Miné-Hattab and M. Smith for their input on data analyses.

